# An Integrated Framework for Genome Analysis Reveals Numerous Previously Unrecognizable Structural Variants in Leukemia Patients’ Samples

**DOI:** 10.1101/563270

**Authors:** Jie Xu, Fan Song, Emily Schleicher, Christopher Pool, Darrin Bann, Max Hennessy, Kathryn Sheldon, Emma Batchelder, Charyguly Annageldiyev, Arati Sharma, Yuanyuan Chang, Alex Hastie, Barbara Miller, David Goldenberg, Shin Mineishi, David Claxton, George-Lucian Moldovan, Feng Yue, James R. Broach

## Abstract

While genomic analysis of tumors has stimulated major advances in cancer diagnosis, prognosis and treatment, current methods fail to identify a large fraction of somatic structural variants in tumors. We have applied a combination of whole genome sequencing and optical genome mapping to a number of adult and pediatric leukemia samples, which revealed in each of these samples a large number of structural variants not recognizable by current tools of genomic analyses. We developed computational methods to determine which of those variants likely arose as somatic mutations. The method identified 97% of the structural variants previously reported by karyotype analysis of these samples and revealed an additional fivefold more such somatic rearrangements. The method identified on average tens of previously unrecognizable inversions and duplications and hundreds of previously unrecognizable insertions and deletions. These structural variants recurrently affected a number of leukemia associated genes as well as cancer driver genes not previously associated with leukemia and genes not previously associated with cancer. A number of variants only affected intergenic regions but caused cis-acting alterations in expression of neighboring genes. Analysis of TCGA data indicates that the status of several of the recurrently mutated genes identified in this study significantly affect survival of AML patients. Our results suggest that current genomic analysis methods fail to identify a majority of structural variants in leukemia samples and this lacunae may hamper diagnostic and prognostic efforts.

## Introduction

Genomic analysis of tumors has stimulated major advances in cancer diagnosis, prognosis and treatment, shifting the focus from morphological and histochemical characterization to consideration of the landscape of driver mutations in the tumor (Vogelstein et al. 2013; Zack et al. 2013; Berger and Mardis 2018). This has been particularly true for leukemia, and especially so for acute myeloid leukemia, in which the spectrum of driver mutations provides a much more rigorous classification of disease subtypes, with a correspondingly more robust prognostic power, than previous histological characterization (Metzeler et al. 2016; Papaemmanuil et al. 2016).

Somatic driver events in a tumor – point mutations, copy number changes and structural variants (SVs) including insertions, deletions, inversions and translocation – are currently identified by some combination of karyotyping, comparative genome hybridization, fluorescence in situ hybridization (FISH), RNA sequencing and genome sequencing of either targeted gene panels, whole exomes or whole genomes (Mardis and Wilson 2009; Alkan et al. 2011; Zack et al. 2013; Wan 2014; Berger and Mardis 2018). However, a recent study interrogating a variety of cancer cell lines using an integrative framework for detecting SVs, consisting essentially of whole genome sequencing, optical genome mapping and chromosome conformation capture, identified a large number of variants that were undetectable by the standard tools for cancer genome analysis (Dixon et al. 2018). Moreover, some of these previously undetected SVs affected cancer relevant genes through their gain or loss or through alteration in expression. In the latter case, gene expression could be reduced by deletion of an associated regulatory domain or activated by fusion of topologically associated domains, bringing an otherwise inactive oncogene in functional proximity to an active enhancer region. This study strongly suggested that non-coding structural variants are underappreciated drivers in cancer genomes. However, since this study investigated only cell lines, it could not differentiate between cancer promoting variants versus variants that arose during establishment and propagation of the cell line itself.

In this study, we have applied a similar integrative framework to identify structural variants in leukemia patients’ primary tumor samples. In particular, we combined both whole genome sequencing and optical genomic mapping to obtain a significantly enhanced view of structural alterations in a dozen different adult and pediatric leukemia samples. In almost all cases, our analysis identified all the structural rearrangements previously determined by standard karyotype analysis. However, our analysis also revealed hundreds of additional structural variants, particularly insertions and deletions but also inversions and translocations, that were not evident from standard genomic analyses. A number of these variants affected tumor associated genes, whose role in prognosis and treatment in the individual cases could not otherwise have been considered. Our work further confirms that the extent of somatic structural variants has not been fully recognized nor effectively integrated into disease assessment. The methods described here may offer a remedy for that shortcoming.

## Results

### Identification of somatic structural variants in leukemia samples

We used a combination of whole genome sequencing and optical mapping to identify structural variants in blood samples from leukemia patients. Patients included seven adult AML cases, two pediatric AML cases, one pediatric T-cell ALL case, one pediatric B-cell ALL case and one adult B-cell lymphoma (Supplemental_Table_S1.pdf). We performed whole genome sequencing on all samples at an average depth of 50X and optical mapping at 100X coverage on a Bionano Genomics Irys or Saphyr optical mapping instrument. For optical mapping, large genomic fragments (>250 kb) are extracted from cells, fluorescently labeled with a site-specific DNA binding protein and then passed through nanochannels of an Irys or Saphyr chip that force the molecules into a strictly linear conformation. After they are linearized and migrate through the nanochannels, molecules are imaged, with the fluorescent tags providing a bar code that allows subsequent assembly of individual molecules into larger contiguous maps, which are compared to a reference genome to identify insertions, deletions and rearrangements.

Data processing to identify structural variants in individual samples is outlined in Figure 1. Whole genome sequence data was mapped to human genome reference hg38 using BWA and then filtered for structural variants by two independent software pipelines, LUMPY and DELLY. Those variants identified by both programs were retained and sorted into subtypes: deletions, insertions, duplications, inversions and intra-and inter-chromosome translocations. Copy number variants were determined by Control FREEC. Structural variants were extracted from optical mapping data using Bionano Genomics Access software. This process yielded in each sample 1500-3000 deletions, more than 2000 insertions, hundreds of inversions and copy number variants and tens of translocations (Table 1).

**Table 1.**
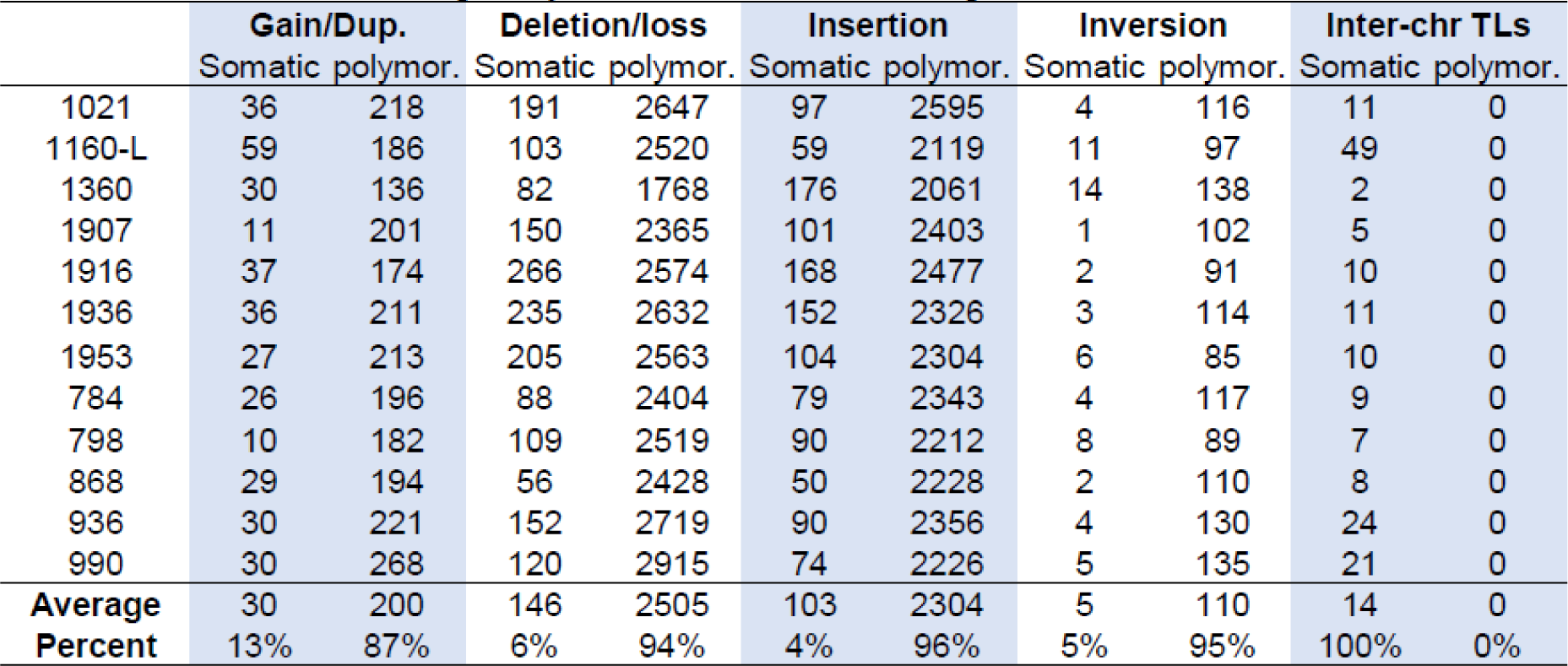
Number of polymorphic and Somatic SVs by combination of OM and WGS

|  | **Gain/Dup.** |  | **Deletion/loss** |  | **Insertion** |  | **Inversion** |  | **Inter-chr TLs** |  |
| --- | --- | --- | --- | --- | --- | --- | --- | --- | --- | --- |
|  | Somatic polymor. |  | Somatic polymor. |  | Somatic polymor. |  | Somatic polymor. |  | Somatic polymor. |  |
| 1021 | 36 | 218 | 191 | 2647 | 97 | 2595 | 4 | 116 | 11 | 0 |
| 1160-L | 59 | 186 | 103 | 2520 | 59 | 2119 | 11 | 97 | 49 | 0 |
| 1360 | 30 | 136 | 82 | 1768 | 176 | 2061 | 14 | 138 | 2 | 0 |
| 1907 | 11 | 201 | 150 | 2365 | 101 | 2403 | 1 | 102 | 5 | 0 |
| 1916 | 37 | 174 | 266 | 2574 | 168 | 2477 | 2 | 91 | 10 | 0 |
| 1936 | 36 | 211 | 235 | 2632 | 152 | 2326 | 3 | 114 | 11 | 0 |
| 1953 | 27 | 213 | 205 | 2563 | 104 | 2304 | 6 | 85 | 10 | 0 |
| 784 | 26 | 196 | 88 | 2404 | 79 | 2343 | 4 | 117 | 9 | 0 |
| 798 | 10 | 182 | 109 | 2519 | 90 | 2212 | 8 | 89 | 7 | 0 |
| 868 | 29 | 194 | 56 | 2428 | 50 | 2228 | 2 | 110 | 8 | 0 |
| 936 | 30 | 221 | 152 | 2719 | 90 | 2356 | 4 | 130 | 24 | 0 |
| 990 | 30 | 268 | 120 | 2915 | 74 | 2226 | 5 | 135 | 21 | 0 |
| **Average** | 30 | 200 | 146 | 2505 | 103 | 2304 | 5 | 110 | 14 | 0 |
| **Percent** | 13% | 87% | 6% | 94% | 4% | 96% | 5% | 95% | 100% | 0% |

**Figure 1.**
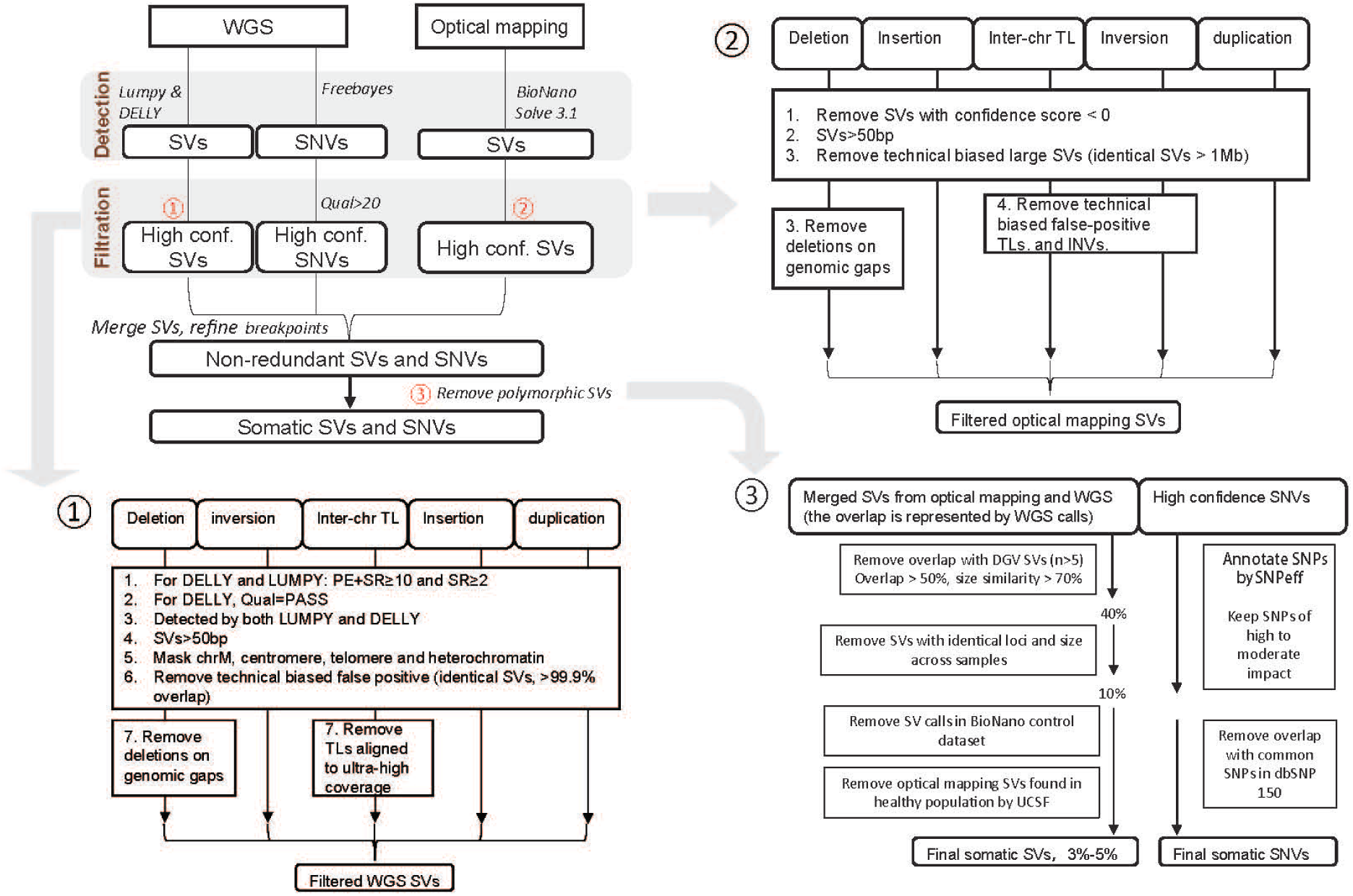
Computational Workflow for Detection of Structural Variants. The computational workflow for extracting structural variants from a combination of whole genome sequencing and optical mapping is diagrammed in the upper left hand figure with details of each of the subroutines provided in the numbered figures. See Materials and Methods for a detailed explanation. Note that step 3 removes likely germ line polymorphisms from the SV calls, reducing the number of original SVs by 95-97% on average.

Determining which of the structural variants arose as somatic mutations versus those that were preexistent in the patient’s germ line would require comparing the list of those present in the leukemia sample to those in the patient’s normal genome. However, since normal tissue is not readily available from most leukemia patients, we developed a computational pipeline to distinguish somatic mutations from germline polymorphisms by filtering the list of variants against various databases of known genomic polymorphisms. We first compared the position and extent of each variant against the Database of Genomic Variants (MacDonald et al. 2014) and removed any variant that significantly overlapped a previously identified variant. We then removed any variant whose start and end point were identical in two or more of our patient samples. Finally, since many of the variants identified by optical mapping could not have been previously revealed by other technologies, we compared our remaining variant list against that obtained from optical mapping of 154 normal individuals in a study recently conducted by Kwok and colleagues (Levy-Sakin et al. 2019), as well as that in Bionano Genomics’ dataset of variants found in normal individuals. As noted in Table 1 and Figure 1, this filtering process significantly reduced the number of variants such that on average only 13% of the initially identified copy number gains and only 5% of initially identified deletions, insertions and inversions were retained as likely somatic variants. In contrast, all of the interchromosomal translocations initially identified were retained as likely somatic events.

We tested the validity of our filtering algorithm in identifying somatic variants in one case in which we were able to obtain normal tissue for the patients. We amplified the small subset of normal T cells from the leukemic blood sample by selective application of growth factors as described in Materials and Methods (Figure 2A-C). We performed optical mapping and whole genome sequencing on this normal T cell population and compared its profile to that of the corresponding leukemia sample to identify somatic variants. We then compared the collection of somatic variants identified by direct comparison to germ line sequences to that obtained by the filtering process described above. As evident from Figure 2C, >95% of the somatic variants identified by our filtering process were not observed in the T cell genome, providing a lower limit for the overall false discovery rate of <0.05. However, due to technical limitations our T cell coverage did not have sufficient depth to identify all the polymorphic structural variants, so we cannot calculate a true false discovery rate. Nonetheless, the results support our filtering pipeline as a convenient, cost effective and likely accurate method for pinpointing somatic variants in leukemia genomes.

**Figure 2.**
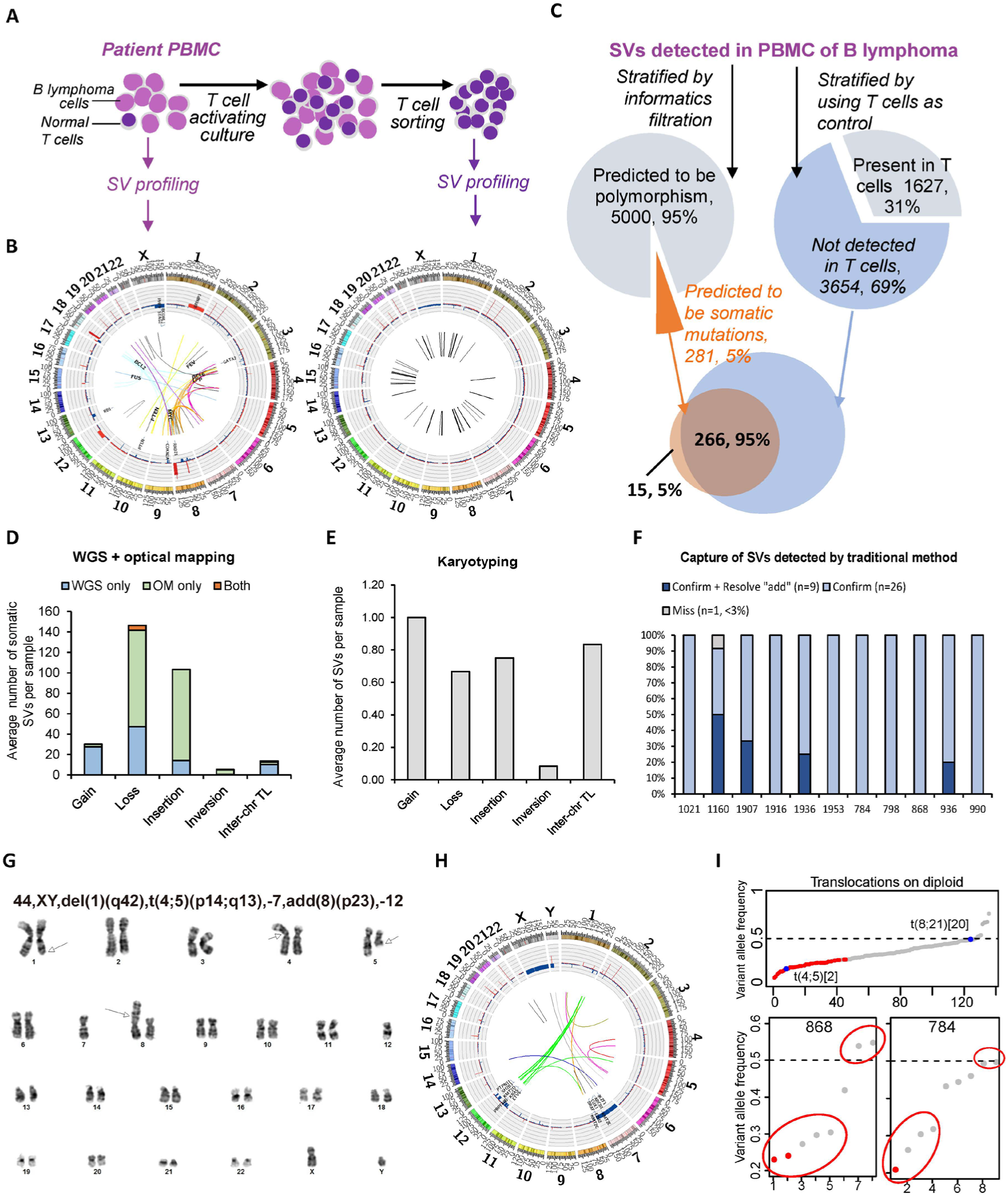
Detection of Structural Variants by WGS+OM versus Karyotyping (A) Workflow for isolating normal T cells from a B lymphoma sample for validating somatic SVs predicted by our bioinformatics pipeline. (B) Circos plots for B lymphoma sample 1160 and the corresponding T cell demonstrate the absence of interchromosomal translocations and the euploid genome in the T cells population. (C)Workflow for determining the false discovery rate for somatic SV prediction by comparison to the T cell control. (D) The average number of different types of somatic SVs over twelve patient samples detected by whole genome sequencing plus optical mapping. Gains include any duplication of genomic sequences greater than 50 bp and losses refer to local or extended elimination of genomic sequences. (E) The average number of SVs detected by karyotype analysis of the twelve patient samples. (F) For each patient sample, the percent of SVs previously identified bykaryotyping that were confirmed by WGS+OM, subdivided into those that were only confirmed (light blue) and those that were confirmed and the source of added material was resolved (dark blue). (G) The karyotype image of sample 936. (H) The circos plot of the structural variants derived from optical mapping and whole genome sequencing of sample 936. (I) Determination of variant allele frequency from WGS data. VAF was calculated as the total number of read spanning the translocation breakpoint divided by the total number of reads spanning the breakpoint plus the total number of read mapping to the intact chromosome at the same site of either one of the participating chromatids. Two translocations identified by karyotyping are indicated by blue dots, corresponding to t(8;21) in sample 784, which was observed in 20/20 karyotype images, and t(4;5) in sample 936, which was observed in 2/20 karyotype images. Lower panels show that VAF separates translocations in each sample into homogenous mutations (upper circle) and sub-clonal mutations (lower circle).

### Comparison of karyotyping, optical mapping and whole genome sequencing

All of the leukemia samples we examined had been previously analyzed by cytological karyotyping as part of the patients’ standard clinical evaluation. Figure 2D and E and Supplemental_Table_S2.pdf presents a comparison of the somatic structural variants identified by each of these methods alone or in combination. As evident, karyotyping revealed only a small fraction of the structural variants present in the sample. Whole genome sequencing alone was adequate for identifying copy number gains and most interchromosomal translocations but failed to identify the majority of insertions and deletions. On the other hand, optical mapping was most effective in identifying inversions, insertions and larger deletions.

As evident from Figure 2D and from previous work (Dixon et al. 2018), WGS and optical mapping provide synergistic data on structural variants. For insertions and deletions, optical mapping picked up larger variants while WGS identified smaller events. In many cases in which WGS failed to flag a variant detected by optical mapping, one or both endpoints lie in an unmappable region of the genome. On the other hand, for some of the variants detected by WGS but unreported by optical mapping, particularly translocations, one side of the variant was too short to encompass at least nine labeling sites, which is the minimum for the mapping software to provide statistically reliable calls. This was particular noticeable in cases of chromothripsis.

Nonetheless, for many of the variants identified by only one of the two methods, the second method does provide confirmation of the validity of the call (Supplemental_Fig_S1.pdf), either by confirming one half of the variant or its reciprocal event. In sum, WGS confirmed 75% of the translocations identified by optical mapping while optical mapping confirmed 39% of the translocations identified by WGS. Finally, we verified a number of the variants uniquely identified by WGS by PCR amplification (data not shown). Accordingly, we are confident that our integrated method reveals a large fraction of structural variants previously unrecognizable.

The combination of whole genome sequencing and optical mapping recovered all but one of 36 genome rearrangement reported by karyotyping (Figure 2F). Moreover, optical mapping plus whole genome sequencing identified 157 interchromosomal translocations in twelve samples, the majority of which were missed by karyotyping. Optical mapping plus sequencing provided a more detailed characterization of translocations than was available from karyotyping. Figure 2G and 2H shows the karyotype image from sample 936 and the corresponding circular genome structure (circos) plot derived from optical mapping and whole genome sequencing. As evident from the circos plot, chromosome 12 had undergone chromothrypsis in this patient’s sample with the majority of the residual fragmented chromosome transposed to chromosome 1. This was not apparent from the karyotype analysis. In a second case, 1021, our analysis documented a three-way reciprocal rearrangement among three separate chromosomes (Figure 3), suggesting that the triple rearrangement occurred as a concerted event. We confirmed the resultant genome org anization by chromosome conformation capture (data not shown). In a number of cases, karyotyping reported that unidentified genetic material had been added to a chromosome without specifying the source of that additional material. In all such cases, our methodology was able not only to identify the source of the exogenous DNA but also to pinpoint the precise junction of the added material (Table 2 and Figure 4). For example, karyotyping indicated additional material on chromosome 12 in patient 1160. Optical mapping identified the extra sequence as arising from chromosome 12 itself, involving an internal inverted duplication of ca. 50 Mb in the middle of the chromosome (Figure 4). This analysis also accounted for the duplicated segment of chromosome 12 identified from our copy number determinations. Circos plots for all the samples analyzed in this study are shown in Supplemental_Figure_S2.pdf.

**Table 2.** Resolution of “Added” sequences

| **Sample** | **Cytogenetics** | **Resolved by OP+WGS** |
| --- | --- | --- |
| BN936 | add(8)(p23), | TL chr8-chr12-2,769,428-96,778,624 |
| BN1160 | add(4)(q31.3) | TL chr3-chr4-148,259,617-156,883,828 |
|  | add(6)(q23) | chromothripsis chr6-chr8-chr18 |
|  | add(9)(p11), | chromothripsis chr9-chr6-chr8 |
|  | add(12)(q22), | Inv chr12-chr12-40,647,772-91,231,824 |
|  | add(14)(q32), | TL chr14-chr18-105,864,250-63,104,165 |
|  | add(15)(p11.1), | TL chr15-chr18-28,135,947-2,823,424 |
| Peds 1907 | add(9)(p13)[7] | TL chr6-chr9-43,687,796-33,130,539 |
| Peds 1936 | add(12)(p13.2)[17] | TL chr12-chr21-11,885,827-34,910,564 |

**Figure 3:**
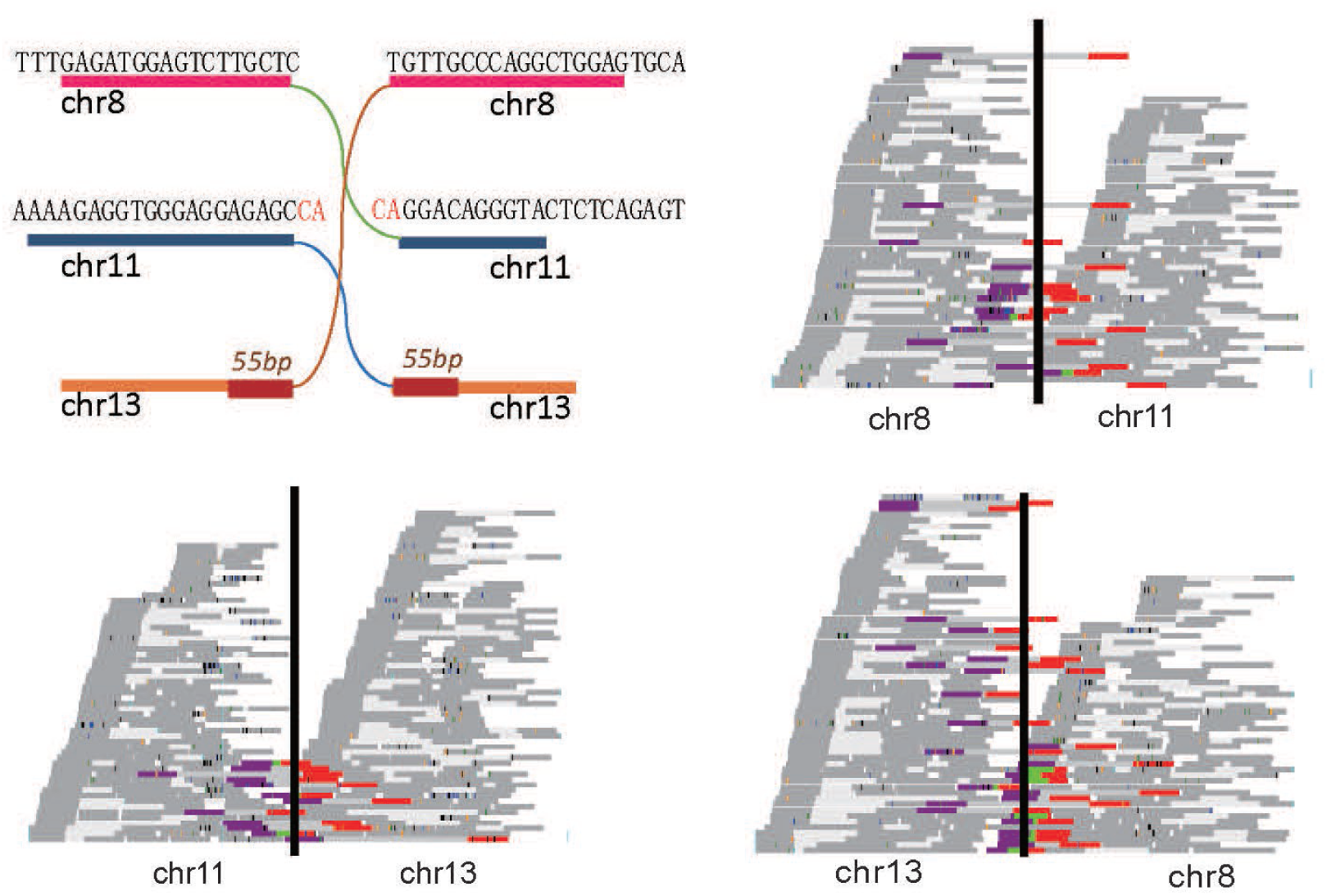
Three-way translocation identified by OM/WGS Diagram of the organization of an unusual three way reciprocal translocation identified in sample 1021 (upper left). The breakpoints of all three chromosomes are precise in that no sequences at the breakpoints are lost and either no sequences duplicated (chr8-chr11), 2 bp duplicated (chr11-chr13) or 55 bp duplicated (chr13-chr8). Pileups of paired-end short read sequences around the breakpoints are shown the other three panels, with each horizontal line representing a fragment on which the dark grey lines represent the sequenced ends of the fragment and the intervening light grey line the inferred unsequenced segment separating them. Reads crossing the boundary are indicated either by dual red/purple colored bars in which the paired ends of a single fragment map to different chromosomes or by tricolored bars in which the green region represents a sequenced segment spanning the junction.

**Figure 4.**
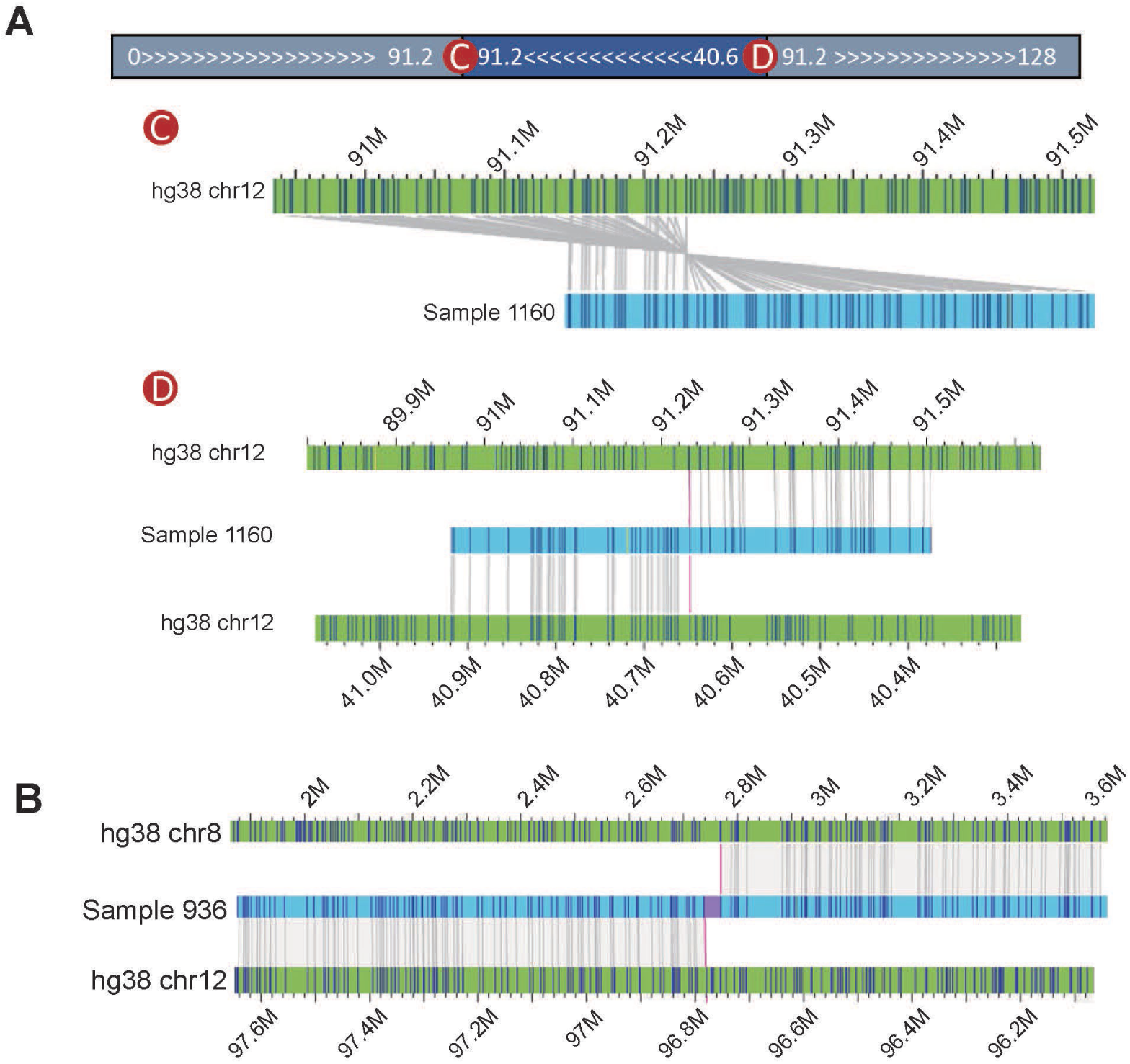
Identification of previously undetermined added chromosomal sequences. Diagram of chromosome 12 derived by optical mapping of patient sample 1160, which had been identified by karyotyping only as chromosome 12 with additional material. Optical mapping indicates that the additional material is a 50 Mb inverted duplication starting at position 91.23 Mb and reconnecting to the remainder of the chromosome at position 40.65 Mb. Below are genome maps showing the hg38 reference chromosome in green with in silico determined labeling sites indicated as dark blue vertical lines, aligned with the chromosomal assemblies around the inversion initiation site at 91.2 Mb (C) and the inversion termination (D) obtained from optical mapping. (B) Genome assembly map in light blue of a chromosome 8/12 fusion obtained by optical mapping of patient sample 936. Detailed mapping of the sample indicates that the chromosome 12 fragment derived from chromothrypsis of one of the chromosome 12 chromatids.

Our method provides information on the relative abundance of a structural variant in a leukemia sample. For instance, calculating the number of reads from WGS that span a translocation breakpoint in relation to the number of reads spanning the intact chromosome at the same site retrieves the allele frequency of that translocation in that sample. For a heterozygous translocation present in 100% of the cells, the allele frequency would be 0.5. As shown in Figure 2I, that is the case for the t(8;21) in patient 784, consistent with the karyotype data indicating that the translocation is present in 20 of 20 images. However, in line with the fact that most somatic structural variants and SNVs are present in subclonal populations in AML cases, we find that the allele frequency of the translocation in most cases is less than 0.5. For instance, we calculated that the allele frequency of t(4;5) in patient 936 is 0.18, consistent with the karyotype report identifying the translocation in 2 of 20 images. Further, in applying this analysis to individual samples, we observe evidence of distinct subclones within the population. As evident in Figure 2I for samples 868 and 784, several translocation cluster at an allele frequency of 0.2-0.3 while other translocations cluster around an allele frequency of 0.5. In sum, our method provides not only the identification of structural variants in patient samples, but also the relative abundance of each variant and evidence of subclones within a sample.

### Functional significance of somatic structural variants

The somatic structural variants identified in our leukemia patient samples overlap those previously associated with leukemia but also affect additional cancer genes. Whole genome sequence analysis revealed single nucleotide mutations or small insertions or deletions in each sample that were previously recognized as driver mutations. Our combined analysis further identified in many samples structural rearrangements in genes previously linked to leukemia and provided a precise definition of nature of the rearrangements. For instance, the precise fusion of BCR-ABL in a 9;22 reciprocal translocation in AML patient 868 emerges from this analysis, information that defines the pathogenicity of the fusion (Melo 1996). In sum, over all twelve patients we identified SVs in thirty-six genes and SNVs in an overlapping set of fifteen genes that were previously implicated as genetic drivers in leukemia, twenty-two of which were mutated in two or more patients (Figure 5A). For example, KMT2C, encoding a histone methyl transferase, is amplified in two patients, deleted in one and carries a point mutation in a fourth (Ruault et al. 2002; Papaemmanuil et al. 2016). Similarly, CEBPA, a CAATT binding transcription factor often mutated in AML (Pulikkan et al. 2017), is affected by SVs in five patients: one case by amplification, one case by translocation, one case by deletion and by single nucleotide mutation in two other patients. Whether these different variants alter the gene function in the same way or affect leukemia progression in different ways in different patients remains to be determined. Only one patient carried an FLT^ITD^ mutation as noted in their clinical reports and as confirmed in our hands by WGS and targeted PCR.

**Figure 5.**
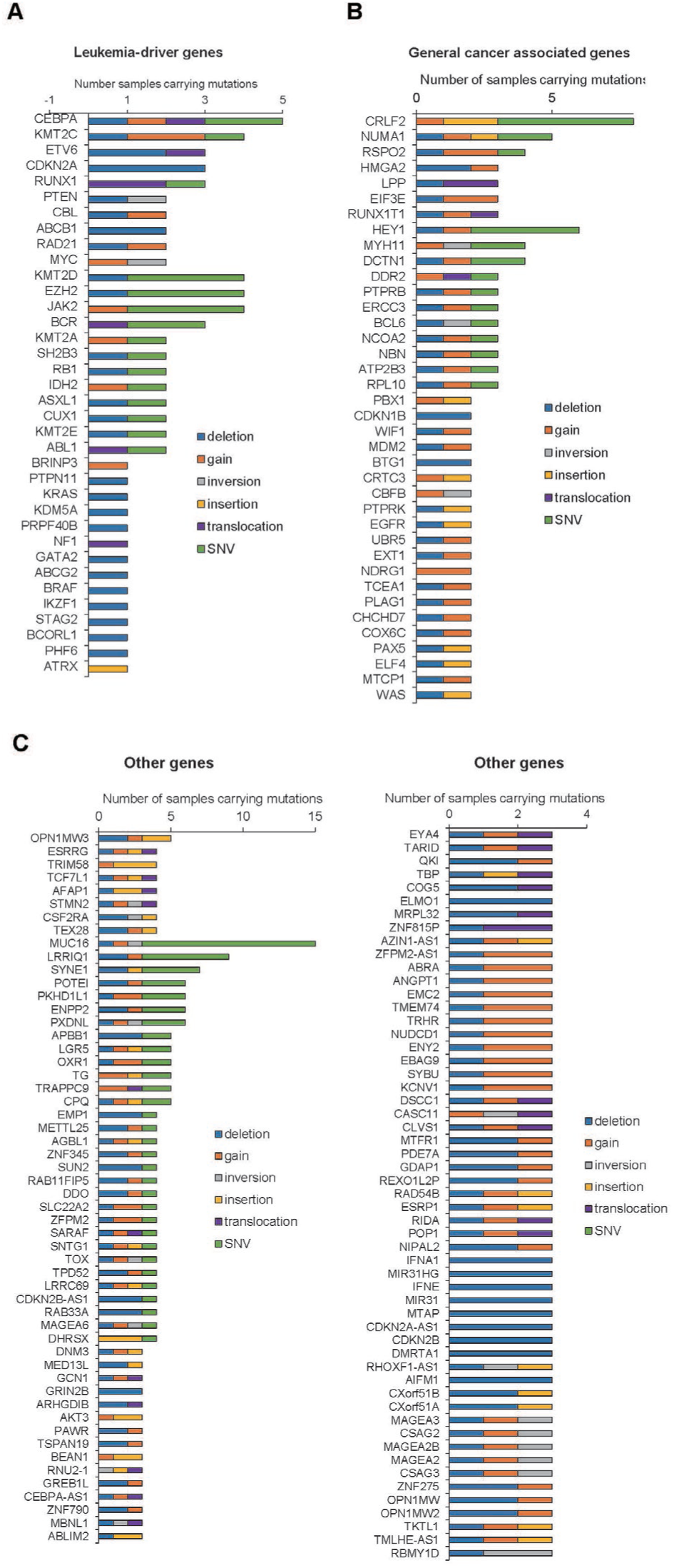
Genes repeatedly disrupted by structural variants in our cohort. The number and type of structural variants identified in our cohort of leukemia patients affecting genes (A) previously associated with AML, (B) previously associated with cancer but not AML, and (C) not previously associated with cancer.

In several cases, we were able to identify loss of tumor suppressor genes that could not be readily detected by conventional methods. In one example shown in Figure 6A and B, a somatic inversion disrupted the PTEN gene on chromosome 10 and a somatic deletion removed the terminal exon of PTEN on its homolog. Neither of these SVs were present in the patient’s clinical report nor identifiable with whole genome sequencing alone. Interestingly, the breakpoints of the inversion correspond to the deletion endpoints in each of two deletions on the homolog. As a second example in Figure 6C and D, BCL6 is disrupted by an inversion on chromosome 3 while its homolog is disrupted by a deletion. As above, neither of these were reported for the patient nor readily evident in the absence of optical mapping.

**Figure 6.**
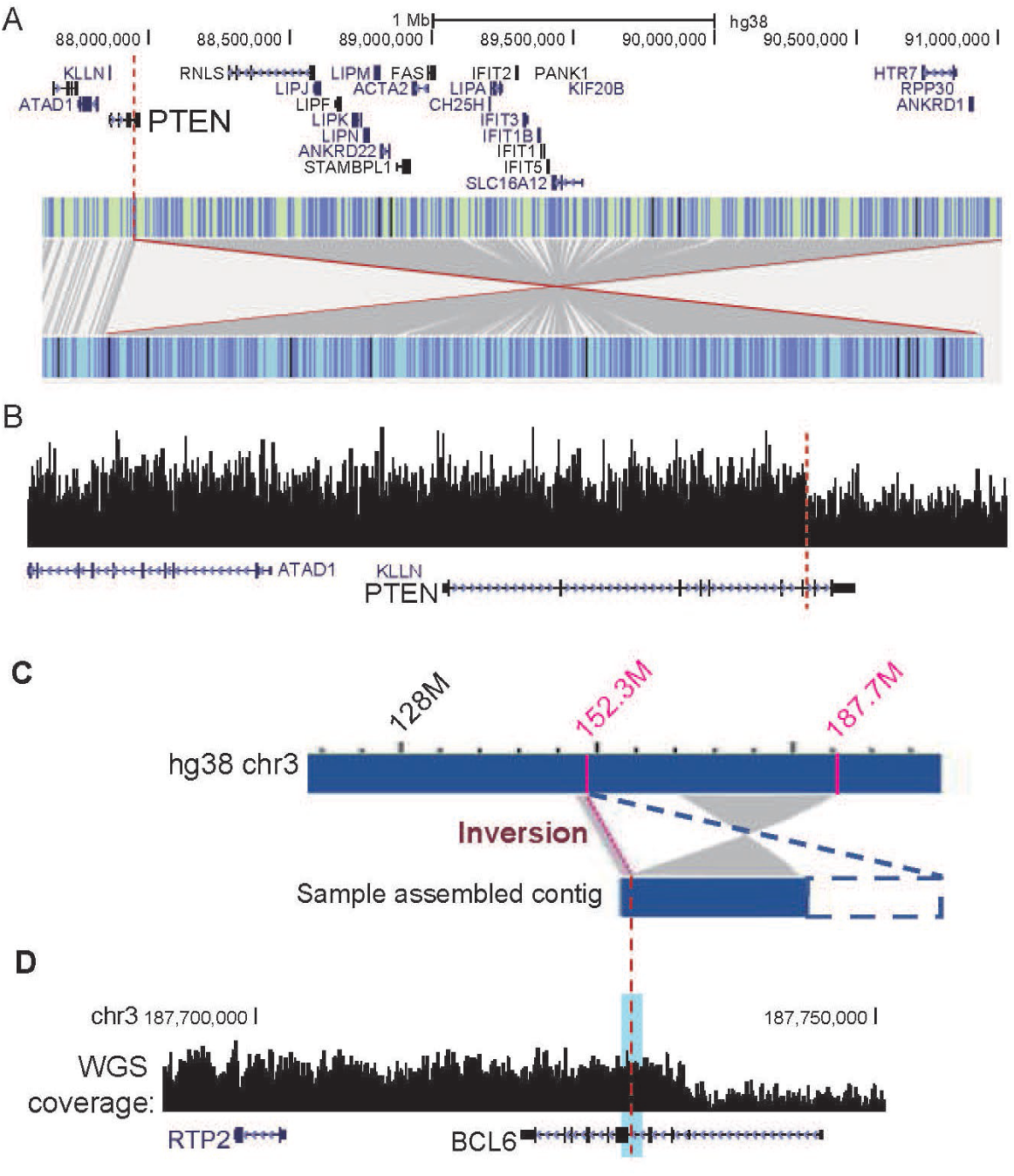
Biallelic disruption of tumor suppressor genes by distinct structural variants (A) Gene map of the region around PTEN scaled and aligned to the in silico generated optical hg38 reference map (light green with blue tic marks indicating the sites of labeling for optical mapping) under which is shown the optical map of patient sample 1160, indicating the position of a 3 Mb inversion, one endpoint of which lies in the PTEN gene. (B) Whole genome sequence read depth over the PTEN region from patient sample 1160 over the scaled and aligned gene map of the region. Dashed red line indicates the start site of a deletion on the chromosome homolog of that in A. (C) Optical map and whole genome sequence coverage (D) from patient sample 1160 positioned over a gene map of the BCL6 locus on which are indicated (dashed red lines) the 35 Mb inversion break point on one homolog and the deletion breakpoint on the other. The reference genome is shown in the top bar, which, due to compression of the tic marks representing labeling sites is solid blue, except for the centromeric region in yellow. WGS coverage level of the BCL6 gene indicates deletion of the last two exons of BCL6 in the second homolog. The current contig contains only one breakpoint of the inversion but does not cover the other breakpoint due to the limited length of the contig. Blue dashed lines represent virtual extension of the contig that is likely the extent of the actual inversion.

We also identified SVs associated with genes previously identified as cancer-associated but not frequently with leukemia (Figure 5B). We found CRLF2 altered in eight patients, twice by insertion, once by amplification and five times by point mutation. CRLF2 encodes a type I cytokine receptor, which along with the IL7 receptor activates the JAK2-STAT pathway, and has been found rearranged in B-cell ALL but not previously in AML (Russell et al. 2009; Harvey et al. 2010; Chiaretti et al. 2016). We also observed alteration in a number of patients of RSPO2, a gene encoding a member of the R-spondin family of proteins that activate WNT signaling. Mutations in RSP02 has been seen in a number of cancers but not previously reported in leukemia (Yoon and Lee 2012; Dong et al. 2017; Wilhelm et al. 2017). As a final example, NUMA1, an essential component in the formation and organization of the mitotic spindle, is altered variously by point mutations, insertion, deletion and amplification. A chromosomal translocation of this gene has been associated with acute promyelocytic leukemia (Cleveland 1995; Wells et al. 1997).

Finally, we observed that a number of genes not previously implicated in cancer were associated with structural variants in multiple leukemia samples (Figure 5C). For instance, eight genes were affected by SVs in at least four of the twelve samples and seventy-five were affected in three patients (Figure 5C). Structural variants affecting the copy number of a particular gene, whether increasing or decreasing, could help drive tumor progression or might simply reflect an adventitious proximity to an actual oncogene or tumor suppressor gene. We have observed several of the later cases for genes near Myc, for instance (Supplemental_Fig_S3B.pdf).

However, many of the novel genes we find repeatedly affected by somatic structural variants are likely contributing themselves to tumor progression.

The AFAP1 gene is repeatedly mutated in our patient samples. The protein encoded by this gene is a Src binding partner that may function as an adaptor protein by linking Src family members and/or other signaling proteins to actin filaments and by mediating Src activation of TGF-β (Gatesman et al. 2004; Qian et al. 2004; Cho et al. 2015). By extracting copy number values from SNP data in a subset of the TCGA AML data, we observed that the AFAP1 coding region is specifically amplified in the AML cohort, while the immediate surrounding region is unamplified (Figure 7B). Moreover, using the TCGA data we found that stratifying the patient population on the basis of AFAP1 expression level provides a statistically significant indicator of patient outcome (Figure 7A). While AFAP1 has not been previously associated with cancer onset or progression, overexpression of the overlapping lncRNA AFAP1-AS1 is correlated with poor prognosis in a variety of cancers, but not leukemia (Ji et al. 2018). However, stratifying TCGA patient outcome data on the basis of AFAP1-AS1 expression or copy number provides no prognostic information (data not shown). Accordingly, we conclude that APAF1 per se plays a role in leukemia progression.

**Figure 7.**
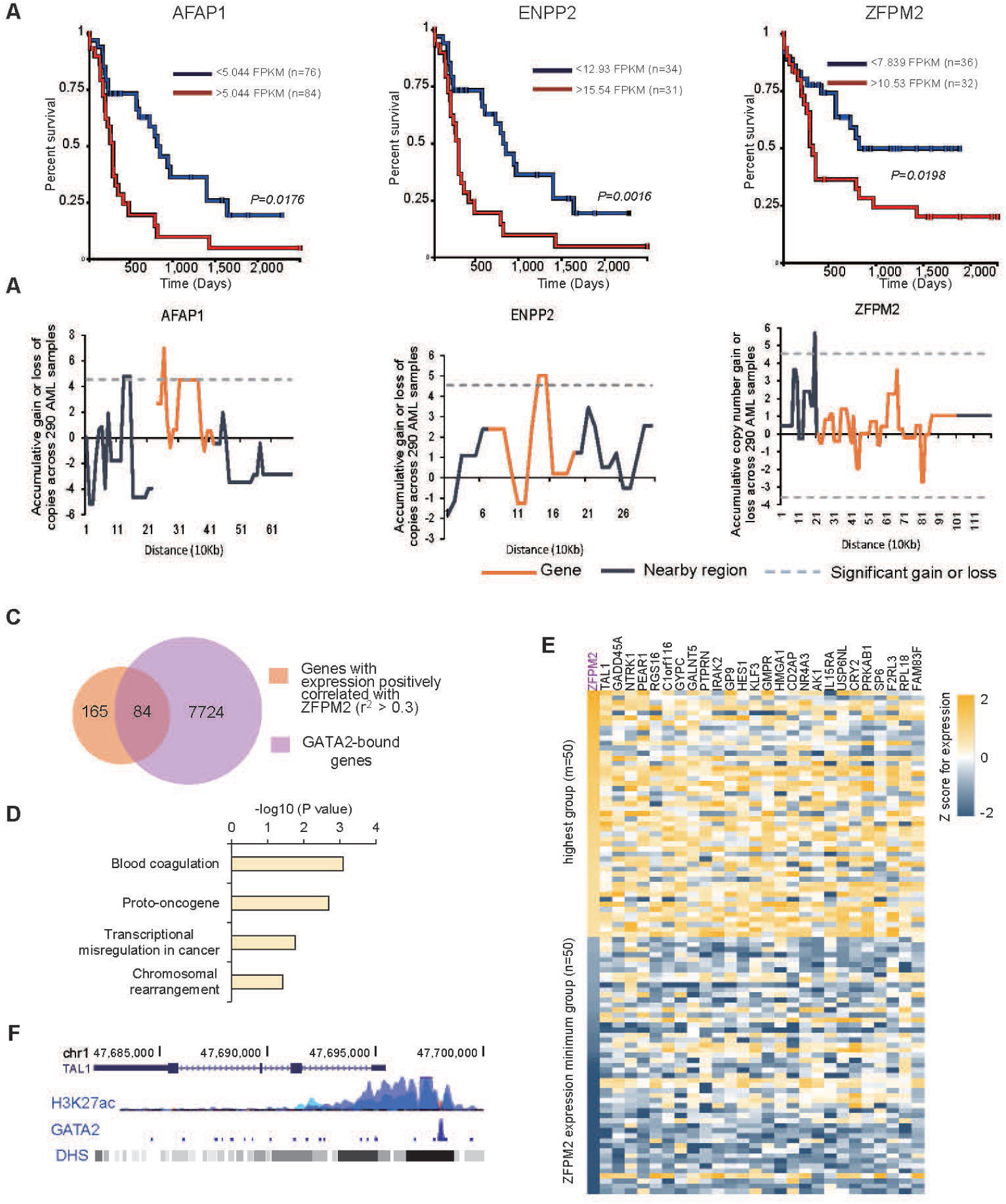
Some genes frequently altered by somatic structural variants affect AML outcomes. Kaplan Meier survival plots of patients in the TCGA cohort stratified on the basis of gene expression for the indicated gene at the thresholds listed for each gene. (B) Genomic copy number alterations of TCGA AML cohorts. Plotted is the z-score for the variation in average copy number in the AML cohort over each 10 kb bin across the gene of interest (orange line) and the adjacent genomic regions (black line). Dotted line indicated the p<0.05 significance threshold. (C) ZFPM2 regulated genes overlap those bound by GATA2. Expression of ZFPM2 exhibits positive correlation with that of 249 genes, 84 of which display binding by GATA2 within the gene body or within 10 kb of the gene in the K562 leukemia cell line. (D) Expression heatmap of a subset of the genes whose expression is correlated and bound by GATA2 for the 50 patients in the AML cohort with the highest ZFPM2 and the 50 with the lowest, sorted by ZFPM2 expression levels. (E) The top four David GO term categories of the 84 genes highlighted in (C). (F) GATA2 binds to the TAL1 promoter. Shown are the genome map of TAL1, the H3K27ac and GATA2 abundance and the DNase hypersensitive sites (DHS) over that region in the K562 leukemia cell line. H3K27ac marks promoter domains.

As a second example, the ENPP2 gene, which encodes the phospholipase autotaxin that catalyzes production of lysophosphatidic acid (Liu et al. 2009), was altered in several patients. Autotaxin is overexpressed in a number of cancers, including breast and ovarian, but has not been associated with clinicopathologic parameters in those or any other cancers (Onallah et al. 2018). In examining the TCGA AML database, we observed that the ENPP2 gene but not the surrounding region is amplified in the cohort and that increased gene expression is significantly associated with worse outcomes (Figure 7A).

As a further example, the zinc finger protein multitype 2 (ZFPM2) gene, also known as friend of GATA-2 (FOG2), encodes a transcriptional cofactor of members of the GATA-binding family that regulates expression of key genes essential for the development of multiple organs (Lu et al. 1999). By interacting with GATA factors, ZFPM2 modulates this regulatory activity, and is known to play important roles in cardiac, gonadal, and pulmonary development. We find that ZFPM2 was affected variously by deletion, duplication and point mutation in four different patients. As above, we interrogated the TCGA AML database, using the genome wide SNP data to determine copy number levels over and around the ZFPM2 gene. We found that the coding region but not the surrounding genome was specifically amplified in patient samples and that high expression of the gene was associated with poor outcomes (Figure 7A). Since ZFPM2 is a transcriptional cofactor, we extracted from TCGA AML data those genes whose expression is correlated with that of ZFPM2 (Figure 7C-D) and showed that those genes significantly overlapped with those bound by GATA2 and were enriched in proto-oncogenes and those associated with transcriptional misregulation in cancer (Figure 7E). As shown in Figure 7F, GATA2 binds to the promoter of one such gene, TAL1, an erythroid differentiation factor (Porcher et al. 2017), which suggests that TAL1 expression may be regulated by ZFPM2.

Finally, a number of genes frequently altered in our cohort but not previously associated with cancer, such as CPQ, COG5, TPD52, AIFM1, RAB33A, ZNF275, TBP and others, provide prognostic information in the TCGA cohort on the basis of their expression levels (Supplemental_Fig_S4.pdf). None of these genes have altered copy number as a consequence of adventitious local amplification or deletion (Supplemental_Fig_S3.pdf). In sum, this study has revealed a number of previously unrecognized structural variants affecting leukemia associated genes as well as recurrent mutations in other cancer associated genes and genes not previously associated with cancer. Outcomes data suggest that some of these newly identified genes could have significant prognostic value.

### Structural variants in non-coding regions affect expression of cancer associated genes

In addition to structural variants that affect the coding region of suspect genes, we observed structural variants near genes but not affecting their coding region. Such structural variants could alter the expression of the adjacent gene by deleting a cis-acting regulatory element such as an enhancer, by duplicating an enhancer element or by fusing the gene to a novel enhancer (Lupianez et al. 2015; Franke et al. 2016). To determine if structural variants in non-coding regions might affect gene regulatory regions, we asked where the endpoints of the different subclasses of structural variants in our patient cohort lay with regard to the boundaries of the approximately 500 AML and general cancer associated genes. We removed from consideration any structural variant that interrupted a gene’s coding region. We then shuffled the genome one hundred times with regard to the cancer related genes and after each shuffle, we calculated how many cancer genes were within a specific range of an SV, and this was calculated for each subclass of SVs. By comparing the distribution of distances from the in silico randomization process to the actual number of genes residing at a specific distance from an SV, we could calculate a z-score reflecting the degree of enrichment of a specific class of SVs lying near a cancer related gene. The results of this analysis, shown in Figure 8A and B, suggests that the endpoints of translocations and deletions are enriched in the 5 Mb region adjacent to cancer related genes, and likely even closer (Supplemental_Fig_S5.pdf). Enhancers are typically found in this region, suggesting that intergenic structural variants in our patient samples might be affecting expression of cancer genes. The cancer genes lying near an SV in each patient sample are listed in Table 3.

**Table 3.**
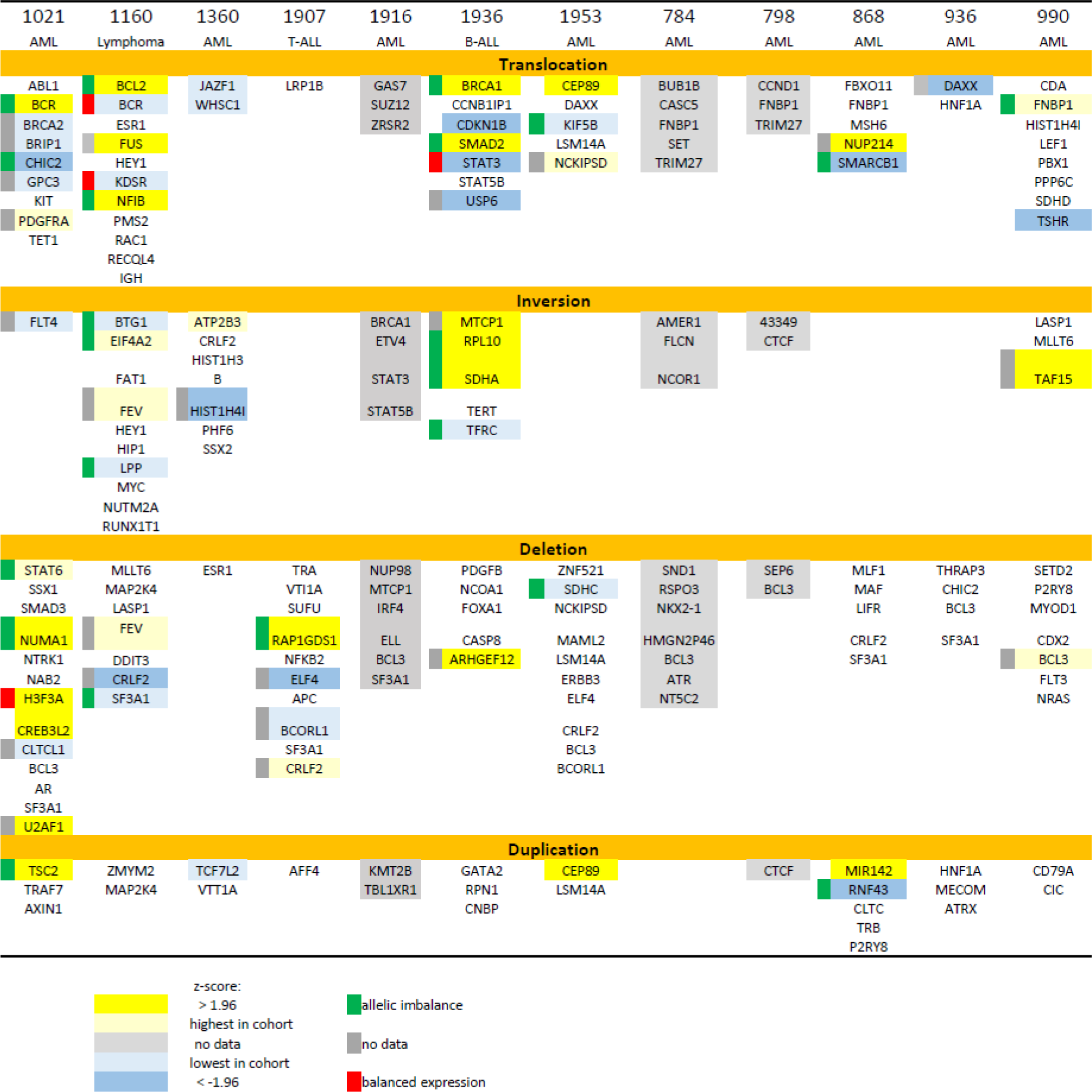
Cancer Genes Adjacent to Structural Variants

**Figure 8.**
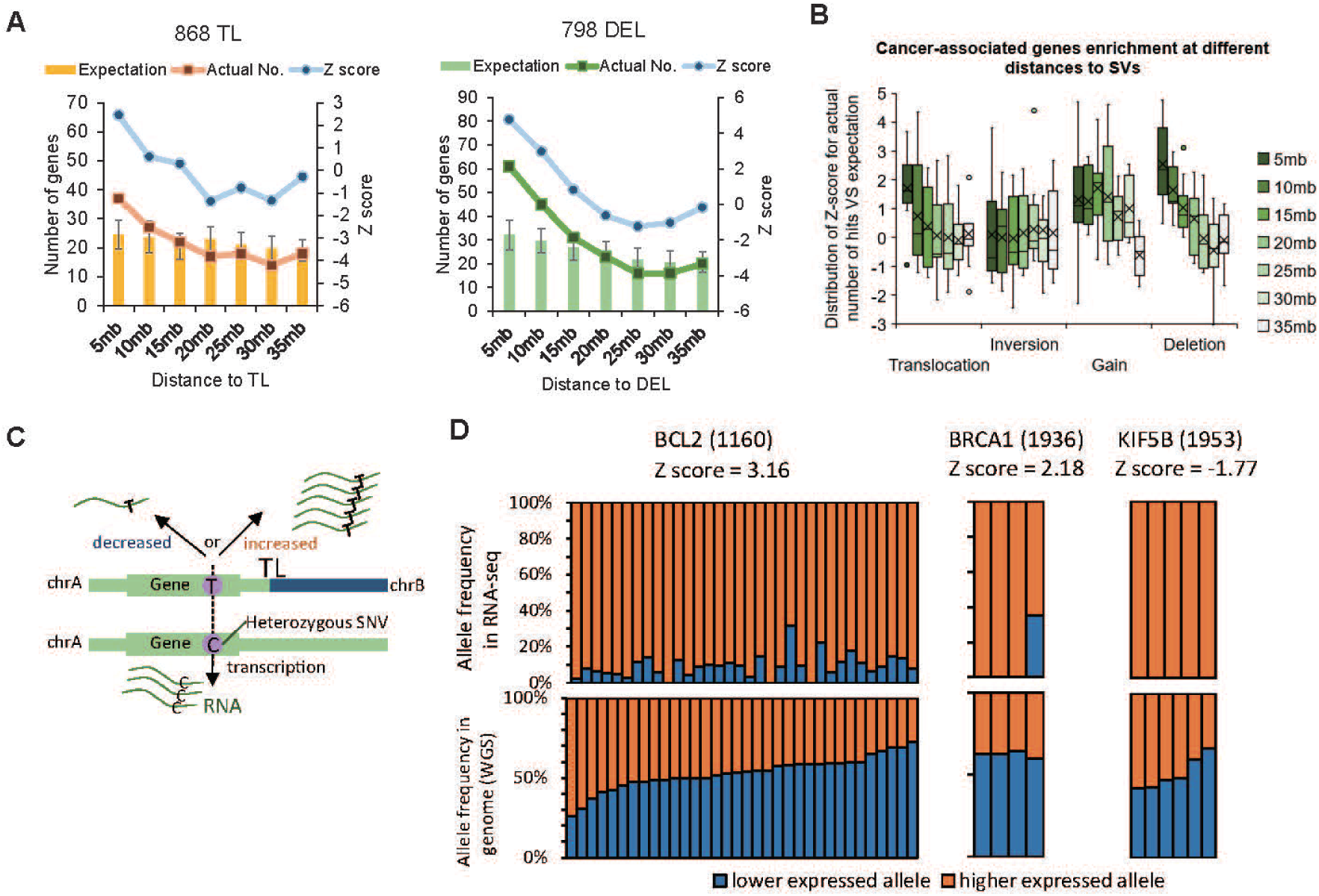
Intergenic SVs affect expression of genes in cis. (A) Likelihood of a translocation or deletion falling near a cancer gene in specific patient samples. The location of any of 500 known cancer genes relative to the closest translocation in sample 868 or deletion in sample 796 was determined and compared to 100 randomized relative positions to calculate a z-score likelihood that a cancer gene would lie in the particular indicated interval. Predicted, actual and z-scores are plotted for each distance interval. (B) Likelihood of intergenic structural variants falling near cancer genes. These calculations for all the samples in our cohort indicate that the translocation and deletion endpoints are enriched in the 5 Mb region upstream of genes, where enhancers are generally located. (C) Schema for determining allelic imbalance of expression of genes lying near intergenic structural variants. (D) Plotted for the indicated genes in the indicated samples are the allelic ratios for heterozygous SNPs across the gene in that sample as determined by whole genome sequencing (lower panel, ordered by increasing ratios) and by RNA sequencing (upper panel).

To test whether SVs alter cancer gene expression in our cohort, we performed whole transcriptome analysis of our leukemia samples by RNA sequencing. We then merged our raw expression data with that from the TCGA study, quantile normalized the merged data set and then determined the average gene expression of all genes. We then assessed whether the expression of a cancer related gene lying near an SV endpoint in our cohort differed significantly from the average expression of that gene over all samples. As evident from the data in Table 3 and Supplemental_Table_S3.pdf, 34% of cancer genes lying near somatic structural variants exhibited significantly altered expression relative to the combined TCGA cohort or were the highest or lowest expressed sample in our cohort. We predominantly observed overexpression of the cancer gene suggesting that the structural variant relocated the gene to a new, stronger enhancer or duplicated a preexisting enhancer. In a few cases, we observed reduced expression of the target gene, an unexpected outcome given the expected heterozygosity of the structural variants. However, in only one of these cases was the structural genes altered in copy number (Supplemental_Fig_S6.pdf), indicating that for all other genes, altered expression was a consequence of perturbation to a cis or trans-acting regulatory element.

To test more directly whether the altered expression of these cancer genes was a consequence of altered cis-regulatory elements, we calculated the allelic bias of the transcripts from that gene in the affected patient. For each gene in question, we identified from whole genome sequencing single nucleotide polymorphisms in the transcribed region that were heterozygous in the relevant patient’s sample and then determined the allelic ratio of those polymorphisms in the RNA transcript sequences in the affected patient sample. If the altered expression were a consequence of the intergenic structural variant acting in cis, then we would expect to observe a significant bias in the RNA transcripts, since the variant should affect only one of the two alleles (Figure 8C). Representative results of that analysis are shown in Figure 8D and summarized in Table 3 and Supplemental_Table_S3.pdf. For a number of genes, this expectation is met. For instance, the increased expression of the BRCA1 gene in patient 1936 comes almost exclusively (3:1 allelic bias) from one allele. Similarly, the reduced expression of KIF5B in patient 1953 results from attenuation of expression from only one allele. In sum, more than one-third of the cancer genes adjacent to intergenic structural variants exhibited significantly altered expression and of those for which allelic bias could be assessed, 85% were expressed predominantly from one allele. These data demonstrate that intergenic structural variants, which are not captured by gene panel or exome sequencing, could play a role in cancer gene expression and their associated role in cancer onset or progression.

## Discussion

This report examining leukemia patient samples as well as a previous report examining cancer cell lines (Dixon et al. 2018) documents that application of an integrative framework of genomic analysis reveals a large number of structural variants unrecognized, and essentially unrecognizable, by conventional genomic analysis, including whole genome sequencing. While whole genome sequencing has been extensively applied to cancer genomics and optical mapping has been sporadically applied to a few individual samples or cell lines (Jaratlerdsiri et al. 2017; Chan et al. 2018), our study suggests that the combination of the two methods recovers twice as many structural variants as revealed by whole genome sequencing alone and our evaluation of the previous cell line study suggest that this combination is adequate to recover the vast majority of SVs. Furthermore, by comparison to datasets of known polymorphic SVs, we could pinpoint those variants that likely arose as somatic alterations. In one case, we were able to confirm the validity of this computational approach by comparing variants identified in the leukemia sample with those of in the patients’ normal genomes. Thus, our combined procedure provides a facile means to identify somatic SVs in leukemia samples. Moreover, our application of this method to a cohort of leukemia patients revealed recurrent alterations whose relevance would not be evident from evaluation of single samples.

Our study identified somatic SVs in a number of genes mutations of which have been previously associated with leukemia. The study also revealed almost one hundred genes recurrently affected in our patient samples, some of which had been implicated in cancers other than leukemia and some of which had not been previously associated with any cancers. The role variants in these genes play in leukemia onset and progression certainly warrants further investigation. In particular, we are quite interested in determining the therapeutic value of targeting those genes altered in various leukemia samples. For instance, ENPP2 overexpression is associated with poor outcomes of AML patients, suggesting that inhibition of the autotaxin phospholipase activity might improve outcomes in a subset of patients. Similarly, somatic alterations of FGFR1 and FGFR2 in several of the clinical samples might suggest that these patients could be candidates for treatment with the FDA approved Ponatinib.

The previous study on SVs in cancer cell lines documented that a number of deletions led to elimination of enhancers or topologically associating domain boundaries, resulting in altered transcription of associated genes (Lupianez et al. 2015; Franke et al. 2016). We have observed similar loss of cis-acting elements in the primary leukemia samples from our patients and have determined that a number of these variants alter expression of the associated gene. Clearly, determining whether down regulation of expression of these genes attenuates proliferative capacity of the associated leukemia cells would be warranted. For instance, we find that, SMAD2, an intermediary in TGF-β signaling (Blank and Karlsson 2015), is upregulated by an intragenic translocation in one of our leukemia samples. SMAD2 has been shown to be upregulated and over activated in CD34^+^ BM progenitors from MDS patients. Moreover, pharmacologic inhibition of the TGF-β pathway in vivo, using a small-molecule inhibitor of the TGF-β receptor, ALK5, alleviates anemia in a mouse model of MDS (Zhou et al. 2008; Bachegowda et al. 2013). Accordingly, determining whether such pharmacologic inhibitors alter the proliferative behavior of those cells could suggest a novel therapeutic approach for select patients with the disease.

Recent studies characterizing the genomic alterations in AML have generated relative consistent classification systems based on the particular spectrum of driver mutations in a sample (Metzeler et al. 2016; Papaemmanuil et al. 2016). These classifications provide fairly robust prognostic power in predicting the likely outcome of individual patients. Our documentation of SVs provides additional information on the genetic alterations in patients and can refine their classification. Whether this additional information enhances the prognostic capability of the existing classification schemes will require additional correlation of our SV data with clinical outcomes. However, we reported here that stratification of patients on the basis of expression or copy number of several genes we found repeatedly mutated in our cohort provided a statistically significant difference in outcomes. This suggests that additional studies of previously underappreciated structural variants may identify additional useful prognostic markers. Furthermore, these studies offer the potential for providing novel targets for therapeutic intervention.

We have recently determined that optical mapping can be effectively applied to solid tumors. We find that DNA isolated from as little as 10 mg of solid tumor can provide sufficiently high molecular weight material for up to 1000X coverage of the tumor genome. Such level of coverage from such minute amounts of material could provide effective identification of somatic structural variants in solid tumors even when the tumor comprises as little as 30% of sample mass. Thus, we expect that the methodology we have exploited here for liquid tumors can be extended effectively to solid tumors as well.

Can the method described previously and in this report provide an effective replacement for cytological karyotyping of tumor samples? Optical genome mapping of a liquid tumor at 100X coverage can currently be performed for approximately $750 (at volume) while current whole genome sequencing costs for 50X coverage approaches $1500 per sample. These coverages are adequate for the level of analysis needed for somatic variant determination necessary for clinical evaluation. Since the computational methods we describe here for identifying somatic versus germ line variants are both highly sensitive and quite precise, it may be possible to forego the need for a parallel germ line determination in order to identify relevant somatic mutations. Thus, the cost of this novel structural variant determination should be commensurate with that of cytological karyotyping, while the level of detection by this new methodology is substantially greater. However, as evident from the results in this study, we do not yet understand the clinical significance of many of the variants identifiable by this methodology. Thus, the ability to provide guidance to clinicians as to the significance of the data that might emerge from this new methodology is currently limited. However, as we expand this technology to many additional clinical samples, our ability to interpret such structural variant data will clearly improve.

## Materials and Methods

### Patient Samples

Bone Marrow (BM) aspirates or Peripheral Blood (PB) samples were obtained from AML patients, after informed consent using protocols under the Penn State Hershey IRB-approved protocol PRAMS Y00-186 or protocol PRAMS 40532. Mononuclear cells (MNCs) were isolated by density gradient separation (Ficol-Paque, GE Healthcare Life Sciences, Pittsburgh, PA) and frozen for later use. Anonymized adult leukemia samples were obtained from the Penn State Hematology/Oncology Biobank. Anonymized pediatric leukemia samples were obtained from the Pediatric Hematology/Oncology Biobank. Patient clinical and demographic data are summarized in Supplemental_Table_S1.pdf.

### Cell culturing

#### T-cell Expansion from Patient PBMCs

T-cell expansion was performed using the Miltenyi T Cell/Activation Expansion Kit (130-091-441) according to the manufacturer’s instructions. T-cell activation beads were prepared prior to thawing patient cells. 100µl of CD2-Biotin, CD3-Biotin, and CD28-Biotin were added to 500µl of anti-Biotin MACSiBead particles. MACSiBead buffer (0.5% human serum albumin and 2mM EDTA in PBS, PH 7.2, 200µl) was added to the mixture and the beads rotated slowly at 4°C for two hours. Activation beads were stored at 4°C until use. PBMCs from the patient were thawed in TEXMACs media with 1% Pen/Strep and counted using an automated cell counter. T-cell activation beads were added to the PBMCs at 25µl/5×10^6^ cells. IL-2 was added 96 hours later at 0.8µl IL-2/ml of media. Cell cultures were monitored and IL-2 media was added as needed. On day 14 post-thaw, additional T-cell activation beads were added to the culture.

#### T-cell Sorting

4.0×10^6^ cells were harvested by centrifugation and resuspended in 2ml PBS to which was added 5µl of CD3-APC (BD Biosciences −340661) and 20µl of CD33-PerCPCy5.5 (BD Biosciences −341640) followed by incubation in the dark for 15 minutes at room temperature. CD3 positive/CD33 negative cells were recovered by sorting on a BD FACS Aria Sorter II.

### Optical Mapping

#### DNA Preparation

5-10×10^6^ PBMCs or 1×10^6^ T cells were incubated at room temperature for 5 minutes on a table top rocker. The cells were then pelleted at 2000 x g for 2 minutes at 4C. The supernatant was then removed and the cells were washed three times with 3ml of 1x PBS. Cells equivalent to 600ng DNA were embedded in 2% Agarose (Bio-rad) and solidified in 4°Cfor 45 minutes. Cells within plugs were lysed in 2ml cell lysis buffer (Bionano Genomics, San Diego, CA) containing 167μl proteinase K (Qiagen, Germantown, MD) for 16 hours at room temperature. An additional Cell Lysis – Proteinase K solution was added to the plugs and incubation continued overnight. RNase A solution (50 μl, Qiagen, Catalog #158924) was added to the mixture and incubated for one hour at 37°C. Plugs were washed four times with Wash Buffer (Bionano Genomics, Part # 20256) at RT with shaking at 180 rpm, followed by five 15 min washes with 10 ml Tris-EDTA, pH 8.0 (TE) each. Plugs were melted at 70°Cfor 2 minutes and equilibrated at 43°Cfor 5 minutes, to which was added 2 μl of 0.5U/ul Agarase (Thermo Fisher, Catalog # EO046) and incubation continued at 43°Cfor 45 min. DNA was transferred to a 0.1um dialysis membrane floating on 15 ml TE in a 6 cm petri dish and incubated for 45 minutes at room temperature. DNA was transferred with a wide-bore tip to a 1.5 ml Eppendorf tube and incubated at room temperature overnight. DNA was slowly mixed with a wide-bore tip and broad range qubit was performed to measure the DNA concentration in each sample.

#### Nickase Labelin

Each DNA sample was split in two and separately nicked with either 30 U of Nb.BspQI (New England Biolabs) in 1× Buffer 3 (Bionano Genomics) or 120 U of Nb.BssSI (New England Biolabs) in 1× NEBuffer 3.1, at 37 °Cfor 2 hours. Nicked DNAs were labeled at 72°Cfor 60 minutes using 15 U Taq DNA Polymerase (New Engand Biolabs) in 1× each Labeling Buffer and Labeling Mix (Bionano Genomics). Repair of nick-labeled DNA was carried out at 37°Cfor 30 minutes in 1× Repair Mix (Bionano Genomics), 0.25× ThermoPol Buffer (New England Biolabs), 1 mM NAD^+^ (New England Biolabs), and 120 U Taq DNA Ligase (New England Biolabs). DNA staining was performed with the final solution containing 1× flow buffer, 1× DTT (Bionano Genomics), and 3 ul DNA stain (Bionano Genomics), in room temperature overnight. Each of the paired samples underwent an average of 7 rounds of data collection on Bionano Irys platform to reach 100X reference coverage. For each round, 160ng prepared DNA was loaded to a Bionano Irys chip that contains two flow-cells, and each round contains 30 cycles of data collection.

#### Direct labeling

750ng of gDNA was mixed with DLE-1 Buffer (Bionano Genomics, Part#20350), DL-Green (Bionano Genomics, Part#20352), and DLE-1 Enzyme (Bionano Genomics, Part#20351) and incubated for 2 hours at 37°Cin a thermocycler. Proteinase K solution was then added to the reaction and incubated for 30 minutes at 50°C. Finally, a DLS-membrane (Bionano Genomics, Part#20358) was placed upon 60uL of DLE-1 buffer in one well of a DLS-microplate (Bionano Genomics, Part#20357). DNA was transferred onto this membrane, incubated at room temperature for one hour, transferred onto another membrane with DLE-1 buffer and incubated for 30 minutes at room temperature. DTT (Bionano Genomics, Part#20354), Flow Buffer (Bionano Genomics, Part#20353), and DNA stain (Bionano Genomics, Part#20356) were added to the DNA in an amber tube and the tube was mixed at 5rpm for 1 hour at room temperature and then stored in the dark overnight at room temperature. Each labeled sample was added to a BioNano Saphyr Chip (Bionano Genomics, Part#20319) and run on the Bionano Saphyr instrument, targeting 100X human genome coverage.

### Library Preparation and DNA Sequencing

#### Whole genome sequencing

DNA was purified from PBMC or blood using DNeasy Blood & Tissue Kit (Qiagen) or QIAsymphony DSP DNA Mini Kit (Qiagen) as described by the manufacturer. Megabase DNA was prepared using DNA Clean and Concentrator (Zymo Research). Covaris fragmentation of samples was performed using the 400 bp target protocol. WGS libraries were prepared according to the KAPA HyperPrep PCR-free Kit (Roche). Illumina NovaSeq S2 150 bp paired-end sequencing was performed to achieve 40X genome coverage.

#### RNA sequencing

Total RNA was extracted from PBMC isolates using Quick-RNA Miniprep Plus Kit (Zymo Research). Libraries were prepared from total RNA following rRNA depletion with KAPA RNA HyperPrep Kit RiboErase according to manufacturer’s instructions (Roche). Illumina NovaSeq 50 bp paired-end sequencing was performed to obtain 50 million raw reads per library.

## Data Analysis

### Variant detection and filtration from WGS results

#### SV and SNV Detection

We used two pipelines to independently identify SVs. The first pipeline uses BWA-MEM (v0.7.15-r1140) to align the paired-end reads to human reference genome GRCh38 (version GCA000001405.015). Duplicated reads were removed by Sambamba (v0.6.6). Reads with mapping quality ≥ 20 were retained for downstream SV calling by Delly (v0.7.7), which reports SVs as deletion, inversion, insertion, tandem duplication or inter-chromosomal translocation. SVs were also independently detected by the Speedseq pipeline (v0.1.2), in which paired-end reads were aligned to the same GRCh38 reference genome with BWA-MEM. Duplicated reads were removed by SAMBLASTER (v0.1.24). SAMBLASTER then extracted discordant read pairs and split reads for downstream SV detection, which was accomplished by Lumpy (v0.2.13) with default parameters. During the SV detection, Delly and Lumpy exclude a list of telomeric, centromeric, and 12 heterochromatic regions provided by the Delly software (https://raw.githubusercontent.com/dellytools/delly/master/excludeTemplates/human.hg38.excl.tsv).

Copy number variants (CNV) were detected by Control-FREEC (v11.0) with the following parameters “breakPointThreshold = 0.8, coefficientOfVariation = 0.062, ploidy = 2”. Control-FREEC normalizes copy numbers for genome GC contents, mappability, and ploidy. Copy number profile for each 50 kb bin of the genome was used for making Circos plots.

SNVs were detected using FreeBayes (version: v0.9.21-19-gc003c1e, included in SpeedSeq pipeline (version: 0.1.2)) with the following parameters “—min-repeat-entropy 1”. Low-quality SNVs (QUAL field < 20) were removed for downstream analysis. SNVs were annotated with SnpEff (version 4.3) using default parameters and filtered for potential protein altering variants (annotated as high/moderate putative impact). This filtered SNV set was then compared against common SNPs (dbSNP150 with allele frequency > 0.01) to keep only potential somatic SNVs.

#### SV filtration and classification

We employed the following criteria to filter SVs detected by WGS: SVs had to be 50 bp or greater, could not map to chromosome Y or the mitochondrial genome and had to be supported by at least 10 reads combining spanning paired-end reads (PE) and split reads (SR) and an additional 2 split reads. SVs calls from Delly and Lumpy were merged to form a consensus call. Merging criteria differed depending on the type of SVs.

Deletions and duplications were merged if the reciprocal overlap (RO) between calls from Delly and Lumpy was greater than 50%. Deletion coordinates determined by Lumpy were used for the merged call set. Insertions with a RO >= 90% were merged and Lumpy coordinates were used. Translocations were merged if both break-point ends mapped within 50bp of each other and if the strand of the break-point ends matched. Final translocation coordinates were based on Lumpy calls. Coordinates for insertions were obtained from Delly since Lumpy does not detect insertions.

We merged deletions detected by LUMPY/DELLY and loss of copies detected by Control-FREEC to form a non-redundant list of “deletions”. Similarly, we merged duplications detected by LUMPY/DELLY and gain of copies detected by Control-FREEC, removed redundant ones, and defined the overlapped ones as “duplications.” For SVs detected by both LUMPY/DELLY and Control-FREEC, we use the breakpoints provided LUMPY/DELLY.

We excluded inter-chromosomal translocations that were also found in a human normal cell line (GM12878) in order to remove likely polymorphisms. WGS data for GM12878 were downloaded from European Nucleotide Archive (Accession number: ERR194147) and analyzed from SVs by the same aforementioned pipelines. We also removed inversions in each patient sample that share a RO ≥ 99.9% with inversions detected in GM12878. We removed inter-chromosomal translocations whose both break-point ends are within 50 bp in any two individuals, since intra-chromosomal translocations that are shared between two individuals at the nearly same location are likely to be polymorphisms or false positive.

### SV detection and filtration from optical mapping results

#### De novo assembly, SV detection and SV classification

We performed de novo assembly of cancer genomes using long optical mapping molecules, from which we identified SVs by comparing the generated cancer genome to the reference genome GRCh38, using software BioNano solve 3.1.1 with RefAligner and pipeline 7196/7224. DNA molecules used for assembly met the following criteria: length >150Kb and spanning at least nine labels, with a signal to noise ratio higher than 2.75 and backbone intensity lower than 0.6. Parameters used for de novo assembly and SV detection are the same as described in the method section in our previous work (Dixon et al. 2018).

Raw SV output comprises deletions, insertions, inversion, duplications and translocations, which include interchromosomal translocations and any intra-chromosomal translocations that are larger than 5Mb. We ran software smap2vcf to convert SV output to VCF format to determine orientation and then separated intra-chromosomal translocations into deletions (5’->3’) and inversions (5’->5’ and/or 3’->3’) according to their orientation.

#### Filtration of detected SVs

We removed all SVs that were smaller than 50bp and all intra-chromosomal SVs with confidence score smaller than 0. We further removed false positive SVs generated due to technical bias such as similar labeling pattern of distinct genomic regions, which results in misalignment and misidentifications of SVs. We also removed large identical SVs (defined as >99.99% overlap) that were found in more than one sample, since identical somatic SVs are unlikely to repeatedly occur in a small collection of samples. We removed deletions overlapping genomic gaps, which represent correction of gap size of the reference genome rather than true deletions. Finally, we generated a list of false-positive translocations and inversions from our previous work and we removed SVs whose breakpoints that are within 500Kb to these previously identified SVs.

#### Integration of SVs from optical mapping and WGS

We integrated SV calls to combine SVs independently identified by both methods into a single call and to represent each SV with breakpoints of highest resolution available. WGS provides SVs breakpoints with base pair resolution, while optical mapping provides only the nearest labeling site to the left and right of the SV (SV interval) instead of its start and end. We therefore set the following criteria for determining whether SVs independently detected by optical mapping and WGS refer to the same event: 1) Deletions, insertions and duplications detected by WGS must overlap at least 50% with the SV interval demarcated by optical mapping and the difference in size predicted by the two methods must be less than 30%. 2) For translocations and inversions, the breakpoint detected by WGS must lie within 500Kb to that detected by optical mapping and the orientation of the SV determined by the two methods should be consistent. If a copy number gain matches duplications found by optical mapping, we specified the SV to be a duplication. All SVs detected by both methods are represented by the breakpoints obtained by WGS.

#### Determination of somatic SV mutations

We used several filtering strategies to distinguish between polymorphic SVs and somatic mutations. First, we compared our deletions, duplications, copy gain and inversion with corresponding SV type in the database of genomic variations (DGV) (MacDonald et al. 2014) with the stipulation that an SV that appeared in at least five individuals in the DGV is a polymorphism. We removed from our somatic mutation list any SV that overlapped at least 50% with a DGV polymorphic SV with less than 30% difference in size. Second, we removed SVs with identical sizes and positions in any two or more of our samples or in the NA12878 cell line (Dixon et al. 2018). Third, we removed SV calls matching any identified in the UCSF optical mapping dataset of polymorphisms (Levy-Sakin et al. 2019). We interpreted an SV detected by optical mapping in three of the 150 normal individuals in the study to be a polymorphism. Fourth, we removed SV calls that match any observed in the BioNano Genomics control dataset.

#### Circos profiling of leukemia genome

Leukemia genome profiles of each samples were generated using Circos (Krzywinski et al. 2009), which includes three tracks: copy number variation genome-wide, deletions and duplications, and inversions and translocations. We used copy number at 50Kb bin size measured by Control-FREEC. The SVs we plotted were the integrated union from WGS and optical mapping. We display genes that are directly overlapping with deletions and copy gains in the outer track. For inversions and translocations, we set a buffer zone of 50Kb to represent to possible position of SV breakpoint detected by optical mapping. We display genes that are overlapping with the possible position of breakpoints of translocations and inversions in the inner track.

#### Comparing SVs to Karyotype

We defined an SV detected by our method as identical to that identified by karyotyping if 1) the position of the SV detected by optical mapping or WGS corresponds to that provided by karyotyping, demarcated by chromosome and the band on the p or q arm; 2) the SV detected by our method is larger than 1Mb, which would be of sufficient size to be detected by karyotyping; and 3) the type of SV is consistent between methods: deletion or copy loss in our method corresponds to “del” or “-” in cytogenetics; inversions correspond to “inv()”; translocations or insertions correspond to “t()”, “der()” or “ins()”; gain of copies or polyploidy correspond to “+”. Complex forms of copy gain such as fragment duplication, inverted duplication or translocated duplications are generally identified as “add” in karyotyping.

#### Identification of frequently disrupted genes

We intersected RefSeq gene exons (GRCh38) with somatic SVs we detected and considered a gene disrupted if 1) part or all of one or more exons overlaps any part of a deletion, loss or gain of copies, or duplications; 2) the breakpoint of an inversion or inter-chromosome translocations lies within the gene; 3) the coding region carries an indel or SNV resulting in nonsense, frameshift or missense mutation or a splicing sequence alteration. Genes inside of an inversion but not interrupted by the breakpoint are not considered disrupted. We divided genes into three exclusive groups based on data from COSMIC (https://cancer.sanger.ac.uk/census): 86 AML driver genes, 534 other general cancer-related genes, and 23631 other genes without clear evidence for association to cancer.

#### Outcomes analysis

For each novel gene frequently disrupted by somatic mutations, we examined whether its copy number or gene expression correlated with disease outcome. Kaplan Meier survival plots were constructed from clinical outcomes data from GDC AML patient cohorts (https://xenabrowser.net/datapages/; https://portal.gdc.cancer.gov/projects/TCGA-LAML). Patients were stratified on the basis of gene expression or gene copy number evenly into two groups, one containing half of the cohorts with above the average expression/copy number of the gene and the other group containing the half of cohorts with below average expression/copy number of gene.

#### Copy number linkage analysis

We obtained copy number variation of TCGA AML cohort (n=290) profiled by SNP array from GDC (https://portal.gdc.cancer.gov/projects/TCGA-LAML), segmented the genome in 10Kb bins and use an in-house pipeline to calculate the average CNV from all patient for each 10Kb bin. We calculated the Z score and the corresponding P value for each bin genome-wide and used that to set a thresholds above or below which represents significant gain or lost across the AML cohort.

#### Simulating distances between SVs and cancer-related genes

Using 86 previously defined AML-driver gene (Metzeler et al. 2016; Papaemmanuil et al. 2016) and 535 additional cancer-related genes from COSMIC (https://cancer.sanger.ac.uk/census), we calculated the number of such genes within a specific distance interval to the nearest SV for each SV subtype in patient samples as well as gene density (genes per Mb). Genes that directly overlap an SV were excluded. We then permuted the distance distribution by fixing the positions of the SVs and randomly distributing the positions of the list of genes and then calculated the number of genes within specific distance intervals to the nearest SV after each individual permutation. The simulations were run one thousand times to generate a distribution of expected number of genes and gene density for each distance interval. We then calculated the Z-score and P value of the actual gene density by comparing to the distribution of expected gene density for each interval.

#### RNA-seq data processing

RNA-seq reads were processed using the ENCODE standard RNA-seq processing pipeline (https://github.com/ENCODE-DCC/long-rna-seq-pipeline). Briefly, raw RNA-seq reads were mapped to human genome reference GRCh38 (version: GRCh38_no_alt_GCA_000001405.15) with STAR (v2.5.3a_modified). Mapped reads were quantified and aggregated at gene level by RSEM (v1.2.31). FPKM values for each gene were used for downstream analysis. To investigate the level of gene expression of our patient samples in general AML populations, we downloaded gene expression for two AML cohorts from TCGA (link). We then performed quantile normalization for FPKM values across patient sample in this study and TCGA cohorts to eliminate batch effects. To quantify the level of gene expression in this study in general AML population, we calculated the Z-score for each gene on the log-transformed FPKM values relative to the average FPKM value for that gene in all samples both across only our leukemia and across our samples plus the TCGA cohort.

#### Allelic gene expression analysis

We processed RNA-seq data bam file with WASP pipeline (van de Geijn et al. 2015) to correct bias towards certain SNV alleles, which can be introduced during mapping. In running WASP, we input SNVs detected from WGS, and WASP outputs a new bam file with bias removed. We then identify SNV on the newly generated bam file using samtools mpileup.

We then pick all the heterozygous SNVs appeared in WGS data, by the criteria that the allele ratio between reference and alternative alleles should be between 0.333 and 3. We examine the allele frequency of the same loci in RNA-seq data and performed chi-square test on the basis of the allele frequency from WGS and RNA-seq. For each gene, we counted the number of significant SNVs, and we calculated the expression percentage contributed by the dominant allele, normalized by the allele frequency in WGS.

## Data Access

Raw and aligned next generation sequencing files have been submitted to the European Genome-Phenome Archive (EGA; https://www.ebi.ac.uk/ega) within study accession EGAS00001003431.

## Acknowledgement

We would like to thank Dr. Pui-Yan Kwok for graciously sharing data on structural variation in normal individuals prior to publication. This work was supported in part by a grant from the St. Baldrick’s foundation (G.L.M.), from the Four Diamonds Children’s Miracle Network (J.R.B.) and from the Pennsylvania Department of Health using Tobacco CURE funds. The Department specifically disclaims responsibility for any analysis, interpretation, or conclusions.

## Author Contributions

Conceptualization, F.Y. and J.R.B.; Methodology, G.L.M., D.C., Y.C. and A.H.; Investigation, J.X., C.P., F.S., E.S., D.B., M.H., K.S. E.B. Y.C. and A.H.; Writing – Original Draft, J.R.B.; Writing – Review & Editing, J.X. and F.Y.; Funding Acquisition, D.G., S.M., G.L.M. and J.R.B.; Resources, D.C., C.A., A.S. and B.M.; Supervision, G.L.M., F.Y. and J.R.B.

